# Percolation Theory Used to Design Biomolecular Condensates

**DOI:** 10.1101/2023.11.04.565634

**Authors:** Weiwei Fan

## Abstract

Biomolecular condensates formed by liquid-liquid phase separation (LLPS) are crucial for various life activities. Critical phenomena are observed during LLPS in cells and *in vitro*, but few studies provide quantitative theoretical explanations for them. In this study, we set up a Bethe network model to simulate percolation, which explains LLPS quantitatively and semi-quantitatively. We designed a condensate system to determine the peptide’s affinity to its target protein. Finally, we found that the artificial condensate can modify the catalytic reaction’s efficiency. Thus, we provide a new perspective on understanding biomolecular condensate assembly and lay the foundation for artificially designing biomolecular condensates.

---

Membrane-less organelles, such as P granule, cajal body, and stress granule^1-5^, are formed by biomolecular condensates, which participate in living activities such as transcript^6-10^, cell division and differentiation^11-14^, and autophagy^15^. The liquid-liquid phase separation (LLPS) is the driving force behind biomolecular condensate assembly ^16^. In 2012, Pilong Li et al. proposed a multivalent interaction theory that biomolecules with multivalence will form LLPS *in vitro*^17^. The condensate states in cells are determined by the LLPS ability of multivalent molecules in them. J. Guillen-Boixet et al. and D. W. Sanders et al. used computer simulation to present the multivalent interactions and free-energy curves of condensates at multiphase stage^18, 19^. B. Gouveia et al. introduced the capillary force theory to describe the wetting phenomenon of condensates in cells^20^.

Percolation theory describes phase separation and pays more attention to the critical phenomena in the phase separation system and the concentration threshold of components in it. Percolation theory has long been used in polymer research^21, 22^, but it is seldom used to study biomolecular condensates. This theory was introduced to computer simulation of LLPS, but its application to experimentation is relatively scarce.

In this study, we set up a percolation network based on Bethe grid and introduce additional complex network states to simulate experimental conditions. We then apply percolation theory to design artificial systems that can promote the reaction catalyzed by the enzyme under LLPS conditions. Our results shed light on the quantitative research of biological LLPS *in vitro*.

Bethe grid is a tree network without circuit, whose critical percolation probability p_c_ is 1/(n-1), where n is the valance of one node in the network. The percolation probability p is the possibility of one node forms a bond to its neighbored node. We defined the network’s penetrated state as: there is a cluster (called penetration cluster) whose nodes occupy all the layers. Based on this definition, the penetration probability p^∞^ is the possibility of at least one penetration cluster forms. In our simulation, we used the frequency of penetration cluster forming represents p^∞^. At last, we visualized the network as a multilayered concentric circles structure with seven largest cluster colored with rainbow colors.

In our simulation study, we set up a Bethe grid, which is a tree-like network without a circuit, and has a critical percolation probability known as p_c_. The value of p_c_ is 1/(n-1), where n is the valence of one node in the network. We determine the percolation probability p as the possibility of one node forming a bond with its neighboring node. We consider the network to be penetrated when there is a cluster, called the penetration cluster, that occupies all the layers of the network. We use the frequency of penetration cluster forming to represent p^∞^. Finally, we visualize the network as a multilayered concentric circles structure with the seven largest clusters depicted in rainbow colors (Figure 1A).

**Figure 1.**
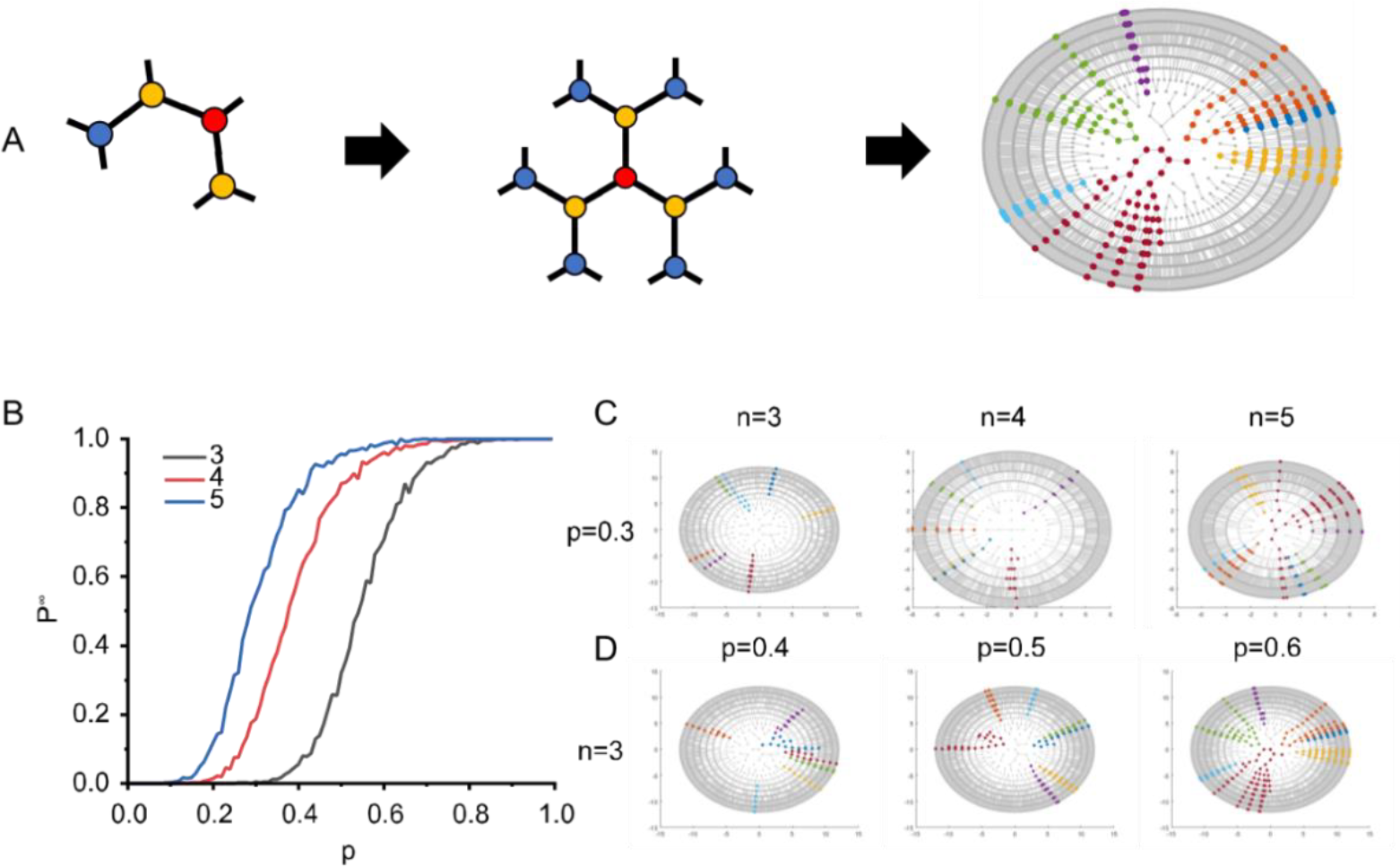
Simulations of the liquid-liquid phase separation on the basic of percolation theory. (A) The visualization of the Bethe grid by the MATLAB. The largest seven clusters are highlighted with rainbow colors. (B) The penetration probability increased as rising of the percolation probability when the valence of the grid is 3, 4, and 5 respectively. (C) The visualizations of the grids with different valence when the percolation probability is 0.3. (D) The visualizations of the grids at different percolation probabilities with the valence of three.

We conducted the simulation with the parameter n=3,4, and 5, at which the theoretical critical percolation probability is 1/2, 1/3, and 1/4, respectively. Our simulation results align with our expectations (Figure 1B), and showed that higher valence n decreased the critical percolation probability p_c_ of the network. We also observed that clusters become larger as the percolation probability p increased (Figure 1C and D). All of these results suggest that the algorithm works as we expected.

Next, we introduced two complex conditions: inhibition and synergy. We randomly deleted bonds formed in the network at different inhibition ratios, which simulated the addition of inhibitors to the LLPS system. As expected, the penetration probability p^∞^ decreased as inhibition ratio increased (Figure S1 A and B). However, obtaining analytic solutions for more complex conditions such as synergy was challenging. To simulate synergy, we modeled the increasing of percolation probability within nodes with decreasing valence of nodes. We observed that when n=3 or 6, penetration was suppressed at synergy ratio of 0.1(Figure S1C). Penetration probability p∞ decreased with synergy ratio rising from 0.1 to 0.3 at n=6. Inversion appeared when synergy gain increased from 10 to 100 (Figure S1D).

Our simulation results indicated that higher valence networks are more robust in complex conditions. We believe that the LLPS system is regulated by many factors and unstable interaction networks may be crushed by them.

To verify the percolation theory for LLPS systems, a “3+2” model was created with adjustable molecular units. Only when the valence of the network is greater than 2, the critical percolation probability is smaller than 1. Due to experimental conditions, interacting units should be separated into different molecules to prevent LLPS formation during protein purification. The ratio of different interaction units should be adjustable to control experimental conditions. The equation for this model is:

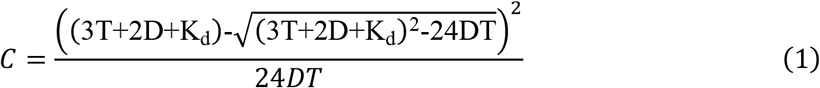

Where T and D represent the concentrations of trivalent and bivalent molecules, respectively. K_d_ represents the dissociation constant, which is the affinity of interaction between protein units and their ligand units, and C is a constant. The numerical simulation results fit well with the theoretical calculation results (Figure S2), but to simplify experiments, the goal was to find the lowest T or D for LLPS. The equation for this was:

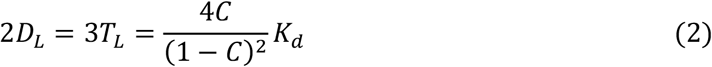

Where T_L_ and D_L_ represent the lowest concentrations of trivalent and bivalent molecules to induce LLPS, respectively.

We used the T2FA-FCP1 complex (PDB ID:1j2x) for this experiment, which consists of three T2FA domains concatenated as one trivalent molecule and two FCP1 peptides concatenated as one bivalent molecule (Figure 2A). The K_d_ of T2FA-FCP1 is 0.5±0.1 μM according to literature^23^. The constant C in equation (1) is 0.5, which is the critical percolation probability of the trivalent Bethe grid. In this “3+2” system, LLPS was observed at concentrations of bivalent and trivalent molecules greater than 10 μM (Figure 2B). The affinity of T2FA and FCP1 was found to be sensitive to the concentration of salt in the buffer. The K_d_ was 10±2 μM at 150 mM NaCl and 24±2 μM at 225 mM NaCl as determined by NMR titration (Figure S3 A and B). We also found that a symmetrical molecule can induce the LLPS of T-T2FA, which expands the scope of applying percolation theory to the interaction of small molecules and proteins (Figure S3C). While there have been few studies that simultaneously determine the affinity and the critical concentration of LLPS, we made every effort and fortunately find the K_d_ of SIM-PRM and the critical concentration of LLPS system with trivalent interaction. The affinities determined by NMR titrations and the critical concentrations of the LLPS were found to be in a proportional function relationship (Figure 2C and Figure S4). These findings in artificial systems provide evidence for the feasibility of applying percolation theory in LLPS research.

**Figure 2.**
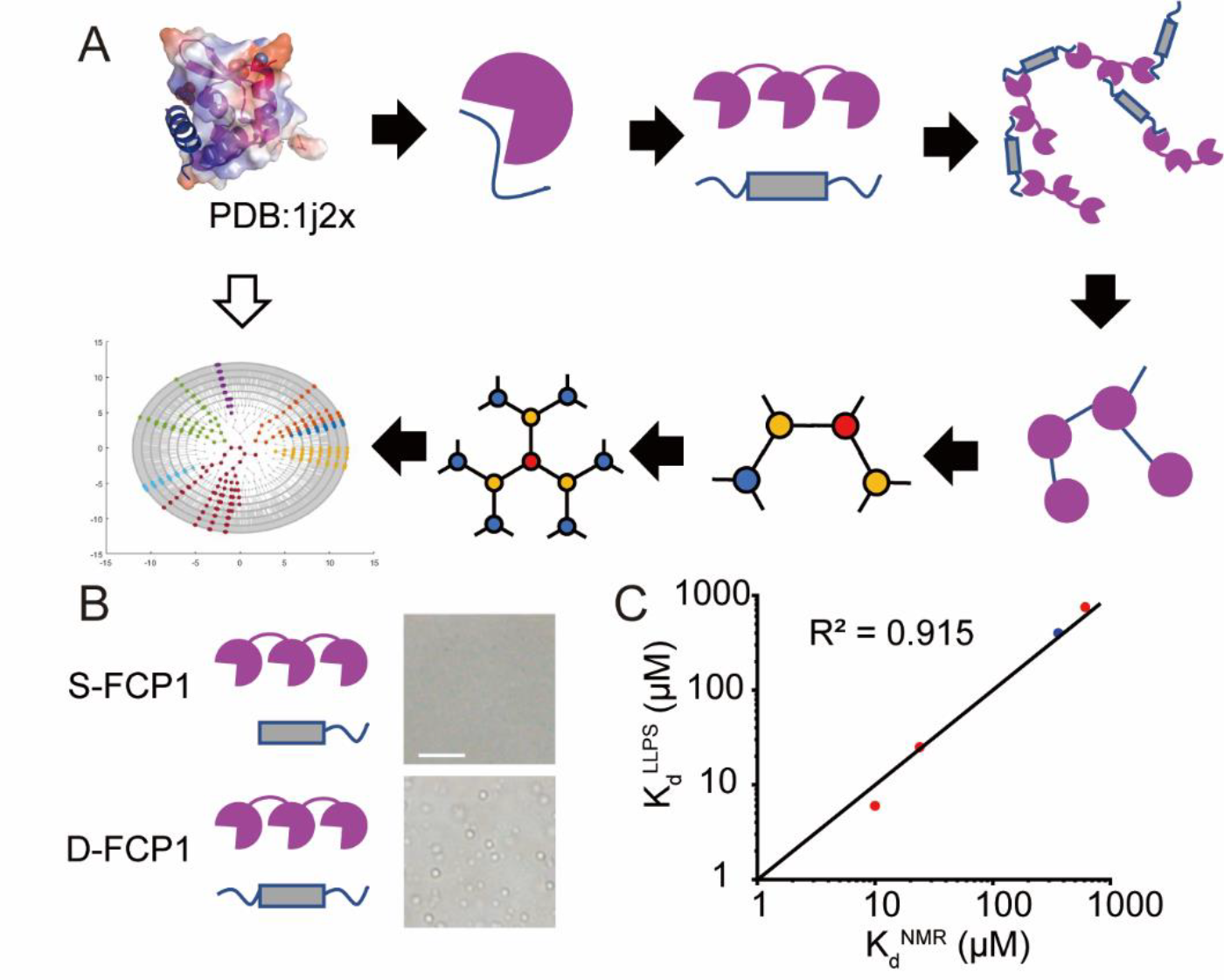
The critical concentration of a specific protein-protein LLPS system is decided by the dissociation constant. (A) The flow chart of constructing the “3+2” system, where the T-T2FA and the D-FCP1 are abstracted as the trivalent node and the bond respectively. (B) Droplets formed in the “3+2” instead of the “3+1” system. T-T2FA: 21 μM (up) and 10 μM (down), S-FCP1: 37 μM, D-FCP1: 15 μM. Bar: 20 μm. (C) Fitting of the dissociation constants determined by the NMR titration and the LLPS.

The T-T2FA protein solution became turbid when the D-FCP1 protein was added, indicating that OD 600 may be a suitable metric to monitor LLPS more conveniently than photography. It was assumed that critical phenomena occur at critical percolation probability, and the observation value v changes as a precipitous S-curve (Figure 3A). The critical phenomenon was observed in the “3+2” system, consistent with the assumed curve (Figure 3B). Furthermore, we created a “3+2+1” model based on the “3+2” model, which introduces inhibitors in the LLPS system (Figure 3C). The equation for this was:

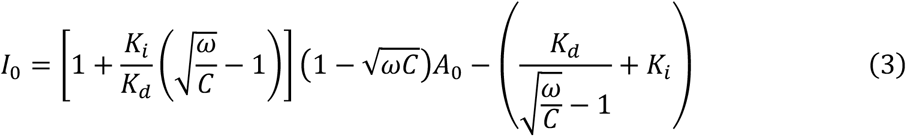

Where I_0_ is critical concentrations of interaction units in inhibitors when concentration of that in trivalent molecules is A_0_. The ω is ratio of interaction units’ concentration in bivalent molecules to that in trivalent molecules. K_d_ and C represent the same thing with these in equation (1) and K_i_ is the dissociation constant of interaction units in trivalent molecules and inhibitors respectively. I_0_ is linearly correlated with A_0_, and it is sensitive to K_d_ (Figure 3 D to F). We used the monovalent FCP1 peptide as an inhibitor to build the 3+2+1 system, with K_i_=K_d_ in equation (3). As expected, I_0_ increased linearly when A_0_ rose, consistent with our theoretical results (Figure 3G).

**Figure 3.**
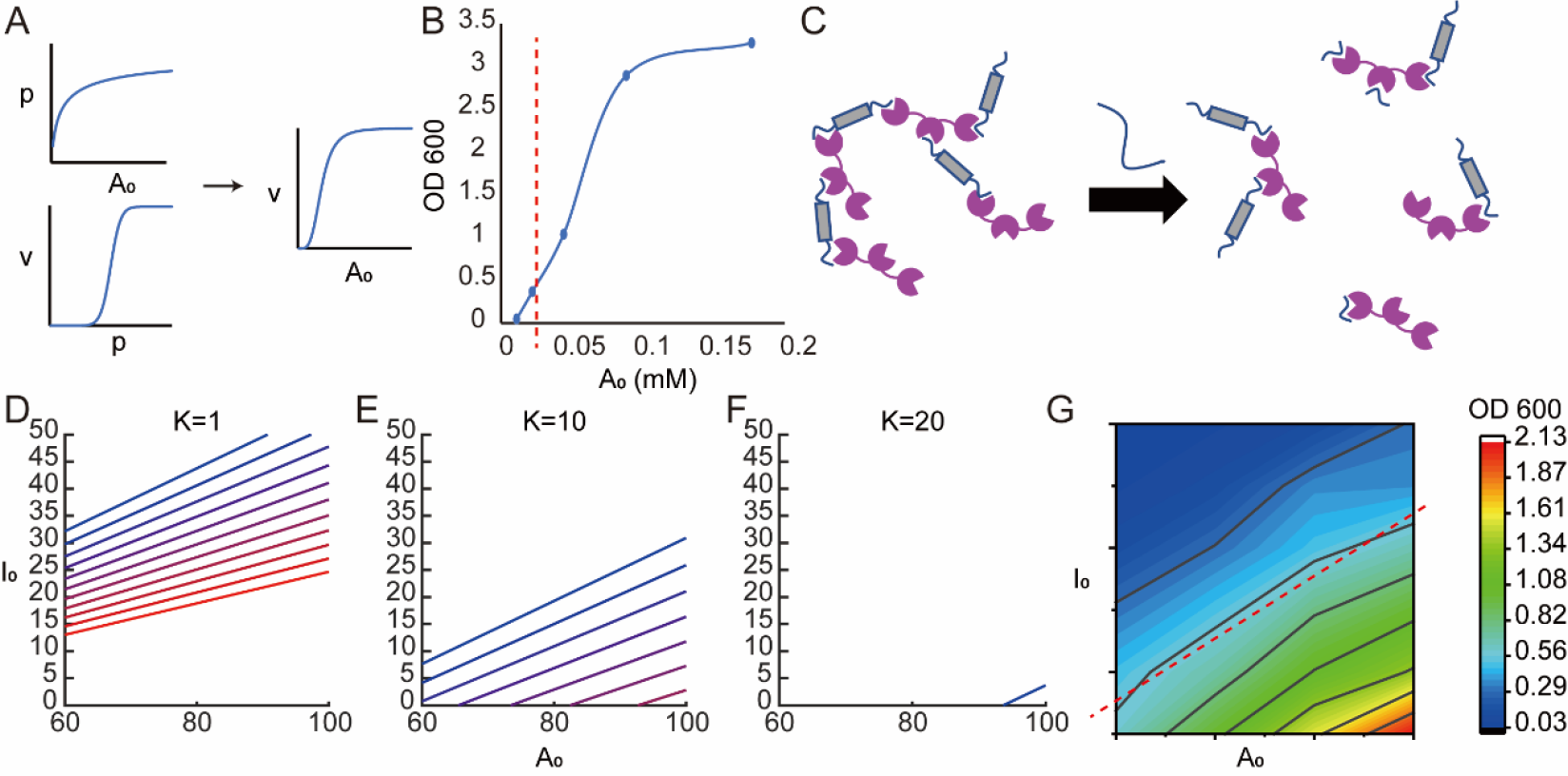
The OD 600 may be a measure of the critical concentration to determine the dissociation constant. (A) v: the observable parameter. A_0_: the concentration of units in multivalent molecules. p: the percolation probability. (B) The curve of OD 600 to the concentration of the T2FA unit. The red dash line represents the theoretical critical concentration. (C) The schematic of the “3+2” system being modified to the “3+2+1” system. (D-F) Curves of I_0_ to A_0_ with C rising from 0.4 (red) to 0.6 (blue), step 0.02, when K is 1, 10, or 20. (G) The OD 600 of the “3+2+1” system at different I_0_ and A_0_. The red dash line represents the theoretical value.

It is believed that proteins form liquid-liquid phase separations (LLPS) to achieve the necessary concentrations for their functions or to alter the efficiency of catalytic reactions. For instance, an enzyme may catalyze a reaction:

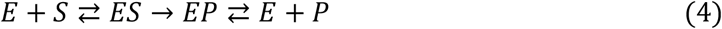

where E, S, and P represent the enzyme, substrate, and product, respectively. ES and EP are intermediate complexes. To simplify the model, it is assumed that the substrate and product share similar affinity to the enzyme and distribution factor inside/outside the droplets. The dissociation constant K and distribution constant K_S_ are also taken into account. Depending on the distribution of the enzyme and substrate/product, there are four different states that are distributed in four different quadrants with K_E_ and K_S_ as axes theoretically (Figure 4A, B). The co-location of the enzyme and the substrate enhances the catalytic reaction, while no co-location suppresses it (Figure 4C). Furthermore, LLPS promotes more binding of the enzyme and the substrate at lower affinity or the concentration of the enzyme (Figure S5A, B).

**Figure 4.**
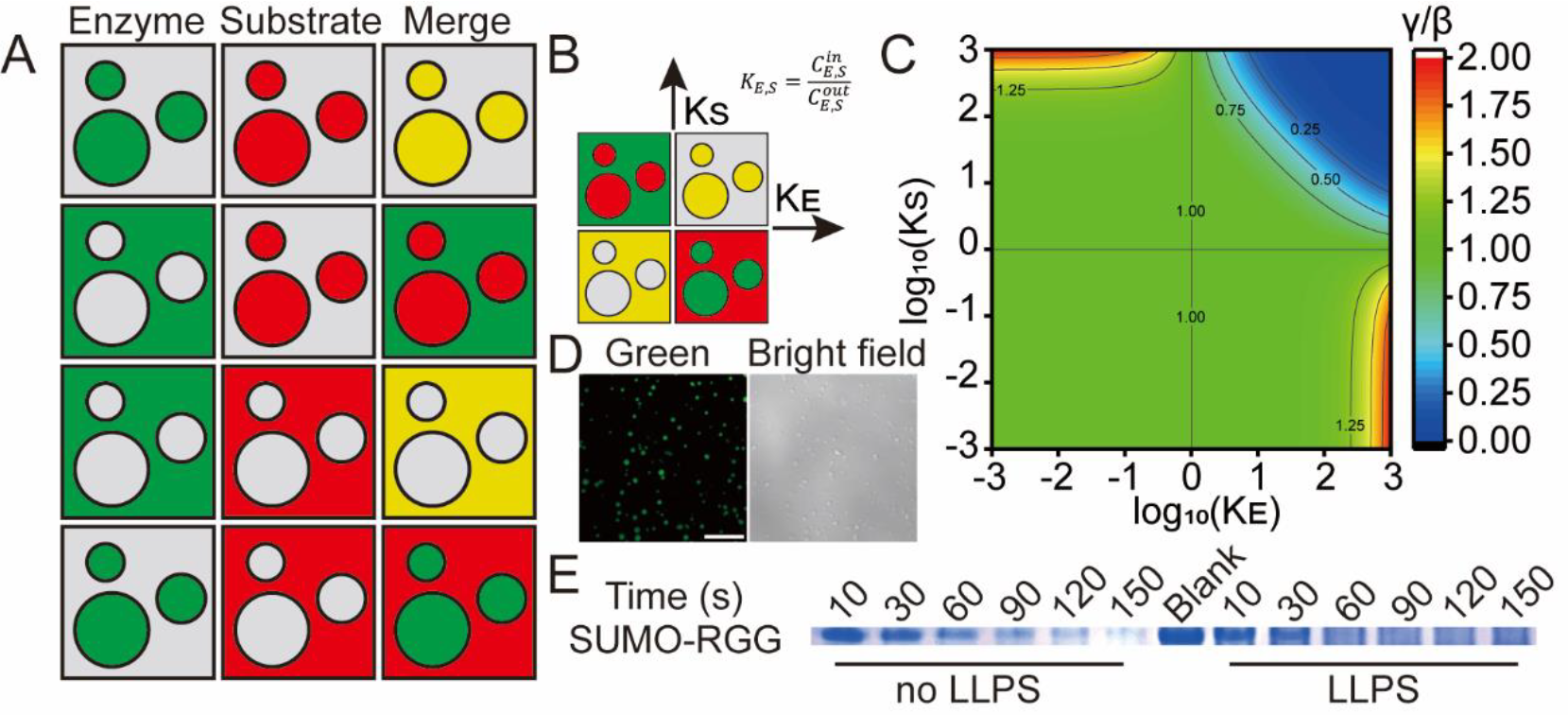
The LLPS accelerates the catalytic reaction when substrates colocalize to the droplets with enzymes. (A) The schematic of enzyme or substrate locations in the LLPS system. (B) K_E, S_ represents the enriching degree of the enzyme and the substrate/product. (C) The lower γ/β represents higher efficiency of catalyzation. (D) the SUMO-RGG was enriched in the droplets assembled by T-T2FA-ULP1 and D-FCP1. Bar: 20 μm. (E) The reaction was accelerated by the LLPS of T-T2FA-ULP1. Blank: SUMO-RGG only, others: SUMO-RGG mixed with T-T2FA-ULP1 and buffer(left) or D-FCP1 (right).

The C-terminal of the SUMO tag is cut by the ULP1, which corresponds to the catalytic reaction illustrated by reaction (4). The substrate protein, SUMO-tagged RGG (SUMO-RGG), is located in the droplets formed by the T-T2FA-tagged ULP1 (T-T2FA-ULP1) and the D-FCP1, and is cut faster as the LLPS is formed (Figure 4D, E). Our results suggest that droplets formed by the LLPS of the artificial “3+2” system can provide a place for biological reactions. This can help mimic those reactions that occur in real cells *in vitro*, giving us a better understanding of how MLOs perform their functions.

Since 2009, there have been two different points of view in the studies on the biological liquid-liquid phase separation: gelatinization and aqueous two-phase system. The former view considers the droplets as a result of crosslinking among biomacromolecules while the latter one views the form of droplets as a free-energy-driven process. However, there is a lack of quantitative research about the biological LLPS that applies either the gelatinization theory or the Flory-Huggins equation. Moreover, it is unclear whether applying the gelatinization theory to the LLPS studies is appropriate. In this study, we introduced the percolation theory parallel to the gelatinization theory, based on which we constructed an LLPS system found to be able to accelerate the catalytic reaction. These results suggest that the percolation theory is appropriate to describe and design biological LLPS systems *in vitro*, which can help explore the physicochemical mechanisms of how MLOs function in cells.

The mathematical expressions of the gelatinization theory and aqueous two-phase system are quite different. The inequality in the gelatinization view is:

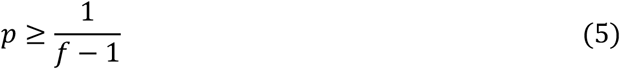

where p and f are the probability and valence of the crosslinks among molecules and the phase transition happens when the inequality holds. On the other hand, in the aqueous two-phase system, the phase separation happens at the binodal line, which is determined by the Flory-Huggins equation:

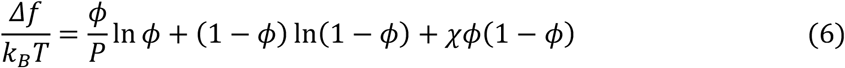

It is almost impossible to determine the required physical quantity in the Flory-Huggins equation in experiments because of the special and fragile characteristics of biomacromolecules. Therefore, the quantitative research is usually based on the gelatinization theory, also called the multivalent interaction theory in biological LLPS studies.

The percolation theory is highly mathematical and is commonly used in the study of solid physics. It shares the same assumptions and similar mathematical expressions with the gelatinization theory but is more solid on the mathematical foundation. Thus, the percolation theory may be a better choice to support the multivalent interaction in the biological LLPS studies. For computer simulation, we demonstrated the uses of the percolation model in a numerical or analytical way (Figure 5A). For a complex system, we used simulation algorithms to predict phase behavior or give explanations for LLPS phenomena qualitatively or semi-quantitatively. Quantitative results were given when the LLPS system is simple, such as few kinds of components and low interaction valence.

**Figure 5.**
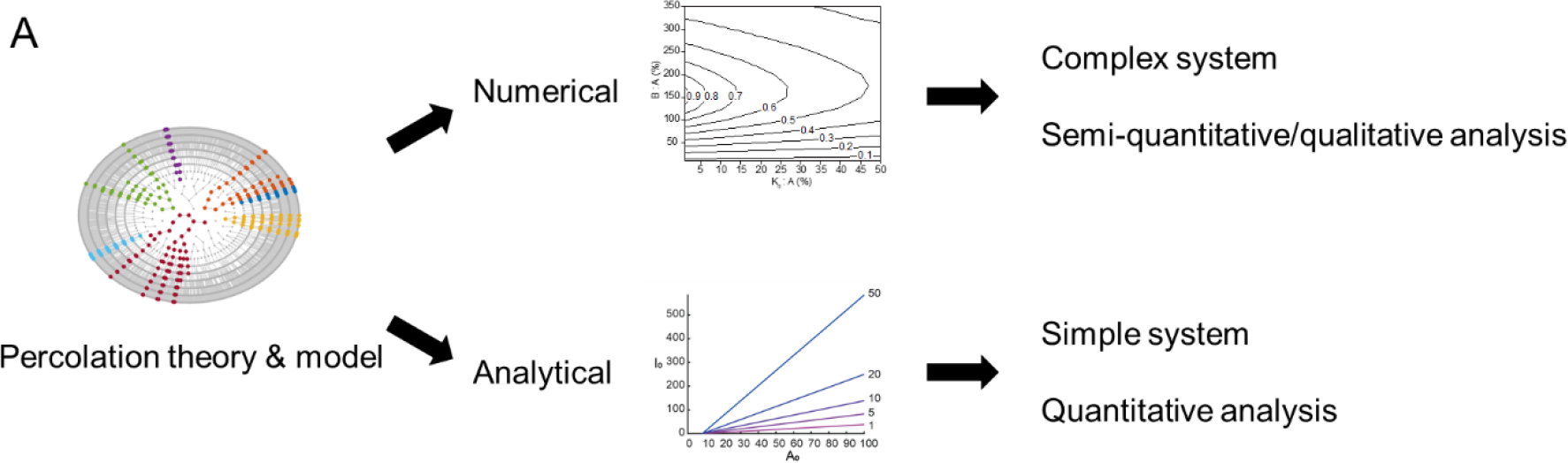
(A) The percolation theory may be used to design LLPS systems which help for studies of bio-condensates i*n vitro*.

For experiments, we designed the “3+2” system that was observed forming droplets *in vitro*. The dissociation constant was found to be proportional to the critical concentration of the LLPS. Moreover, the OD 600 was proved useful in both the “3+2” and the “3+2+1” system, simplifying the monitoring of LLPS. Finally, we raised a model of how the LLPS affects the biochemical reactions and studied the activity of the enzyme in our artificial LLPS system.

In conclusion, we verified the feasibility of applying percolation to biological LLPS research experimentally, which may provide a new perspective and methodology for biomolecular condensates and MLOs studies.

## ACKNOWLEDGMENT

Thanks for the supporting from BioNMR Lab (http://bionmr.ustc.edu.cn/index.html) and Linge Li. Also, thanks for the supporting from Yunge Lou.

## Supplementary Information of

### Materials and methods

#### Cloning, protein expression, and purification

G3BP1 RGG (residues 416-466) were PCR-amplified from a human brain cDNA library and were then cloned into the pET-28a vector. with the SUMO tag. The constructs were then transformed into *Escherichia coli* BL21.

Proteins were overexpressed at 37 °C after induction with 0.5 mM IPTG for 6 hours. Bacterial cells were harvested by centrifugation followed by resuspension in lysis buffer (2 M NaCl and 20 mM NaH_2_PO_4_, pH 6.5). After sonication, lysates were centrifuged, and the supernatants were loaded onto nickel affinity columns (QIAGEN, Shanghai, China) preequilibrated with binding buffer (same composition as the lysis buffer). A washing buffer (binding buffer with the addition of 30 mM imidazole) was used to remove impurities, and proteins were then eluted (binding buffer with 500 mM imidazole). Proteins were further purified by size exclusion chromatography using a HiLoad 16/600 Superdex 75 column (GE Healthcare, Shanghai, China).

The T-T2FA (residues 416-466), D-FCP1 (residues 416-466) and T-T2FA-ULP1 were synthesized by Tsingke Biotechnology Co., Ltd., Nanjing, China). The GHR (residues 416-466) barrier of the D-FCP1 was artificially constructed to reduce the synergy effect. The proteins were expressed and purified as described above.

For NMR experiments, bacteria were cultured in a minimal media (24g NaH_2_PO_4_, 5g NaOH, 0.5g NH_4_Cl, and 2.5g glucose in 1 L H2O), where NH_4_Cl and/or 2.5g glucose was isotope labeled. The expression and purification of proteins were described above.

#### NMR spectroscopy

Proteins were dialyzed against PBS buffer (150 or 225 mM NaCl and 20 mM NaH_2_PO_4_, pH 7.5). The data were processed and analyzed by NMRPipe and SPARKY. The ^15^N-labeled proteins (0.06 mM) were used in the acquisition of 2D ^1^H-^15^N HSQC spectra on an Agilent 500 MHz or 700 MHz system at a series of ligand: protein molar ratios. The dose-dependent chemical shift perturbation was best fitted, assuming a 1:1 binding mode, with Origin. The peptide FCP1 was synthesized by Tsingke Biotechnology Co., Ltd. (Nanjing, China).

#### The derivation of formulas

There is an equilibrium in the “3+2” system:

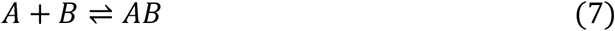

where A and B represent the single unit of the trivalent and bivalent molecules respectively, while AB represents the complex of these two units. There are equations:

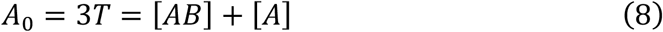

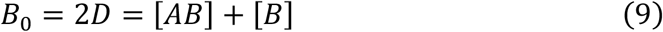

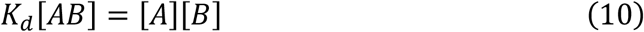

where the [A], [B], and [AB] represent concentrations of the free A, free B, and complex AB. The A_0_, T, B_0_ and D are the total concentrations of the unit A, trivalent molecule, unit B and bivalent molecule regardless of the binding or free states. K_d_ is the dissociation constant of the complex AB. Another equation is often used that:

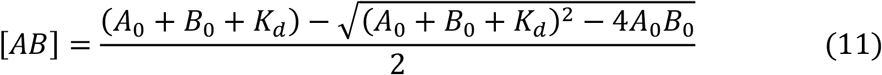

According to the percolation theory, we should calculate the binding probability of two trivalent molecules, which bridged by the bivalent molecule. There are three states of the bivalent molecules: totally free, binding at one side, binding at both sides, whose concentrations are represented by 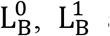 and 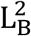. Two get these concentrations, we should calculate the probability of the unit B binding to the unit A, credited as P_B_:

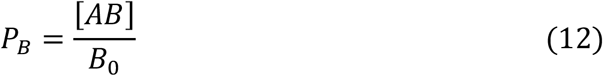

The 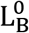, 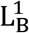 and 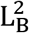 can be calculated by the classic model of probability:

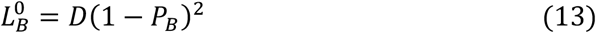

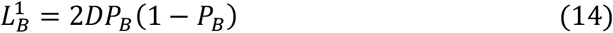

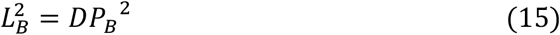

The probability that both sides of the bivalent molecule bind to the same trivalent molecule can be ignored as the huge quantity of molecules in the system. As a result, the probability of linking two trivalent molecules, or called the percolation probability p is calculated as:

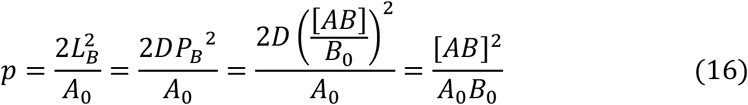

Finally, we can get the equation (1) when p equal to the critical percolation probability C, which is a constant.

To get the minimum of A_0_ or B_0_ when p=C, we should find the derivative of B_0_ to A_0_. Firstly, we modify equation (16) as:

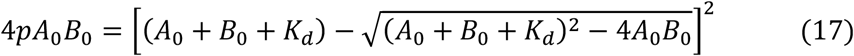

then we expand the right side of the equal-sign and get:

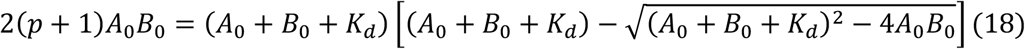

Then we get:

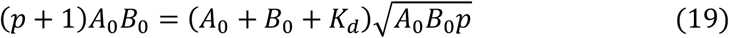

what’s also equivalent to:

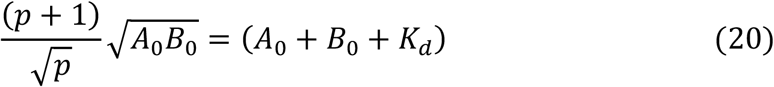

After the differential of both sides, we get

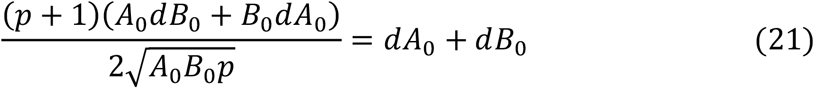

Which means:

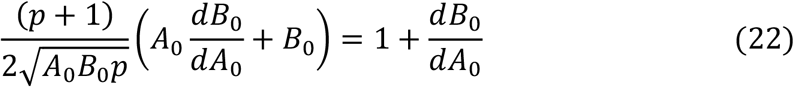

Let 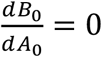, we get:

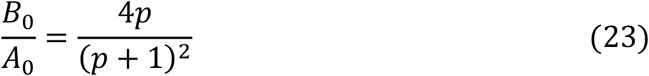

by which, we solve the B_0_ in the equation (20) and get

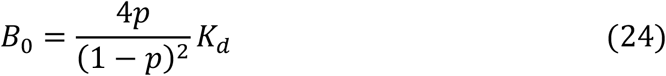

Finally, we get the equation (2) as the curve of B_0_ to A_0_ is symmetrical to the line B_0_=A_0_.

There is another equilibrium in the “3+2+1” system:

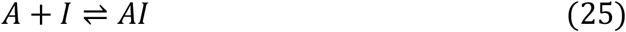

where the I represents the inhibitor molecules and AI represents the complex of A with its inhibitor. The inhibitor changes the equations (8)-(10) as:

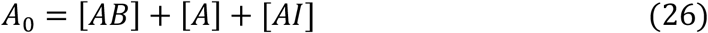

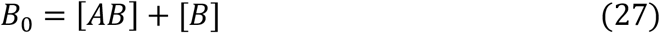

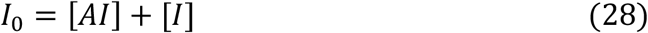

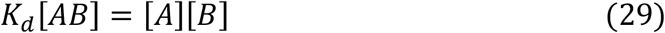

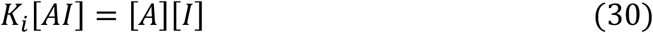

Where I_0_, [I], and [AI] represent the concentration of total inhibitors, free inhibitors, and the complex AI with the dissociation constant K_i_. From equation (28) and (30), we know:

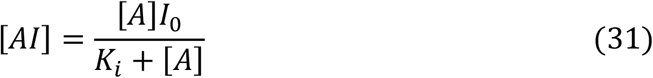

So,

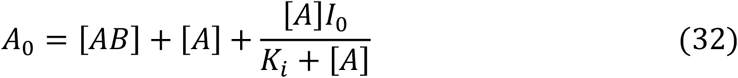

Multiply equation (27) and (32), we get:

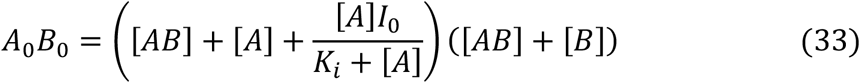

As

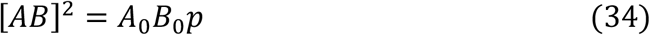

We get:

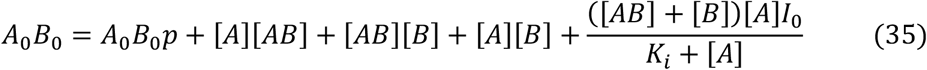

Which equivalent to:

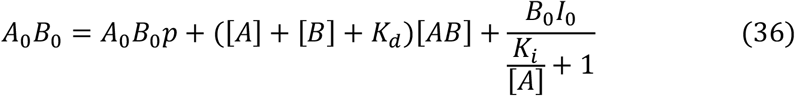

As

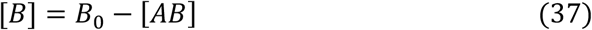

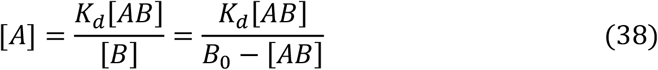

We then get:

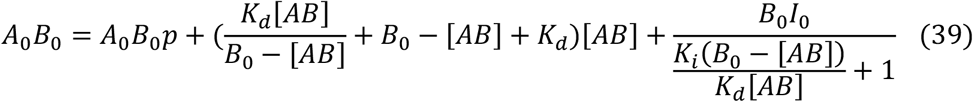

According to the equation (34), we can replace the [AB] and simplify equation (39) as:

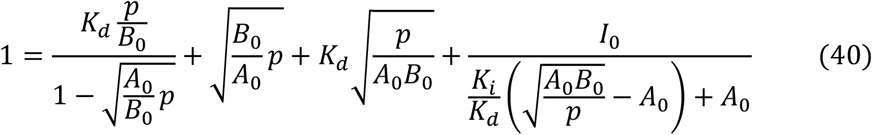

We define the ω as:

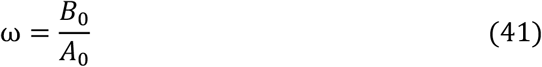

So:

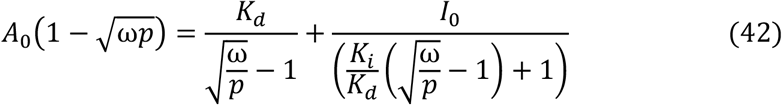

Solve I_0_ when ω is fixed and p is the critical percolation probability C. Finally, we will get the equation (3).

We simplified the reaction (4) system under the LLPS condition as is described in the main body. We assumed the higher concentration of ES and EP represents the higher efficiency of the catalyzation. then we get the equations:

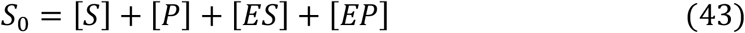

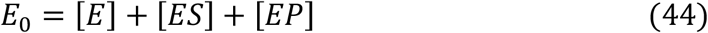

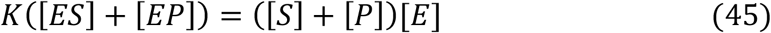

Let

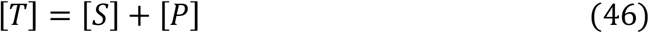

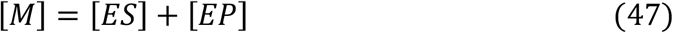

We get:

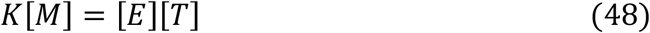

There are some equations describing the LLPS process:

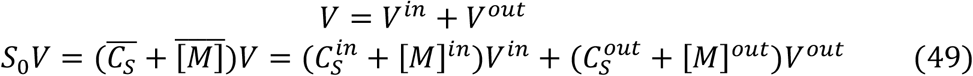

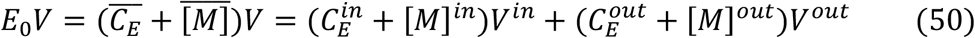

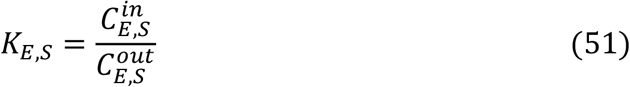

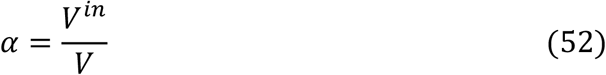

There are also some equations:

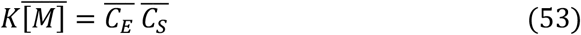

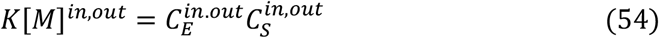

We define:

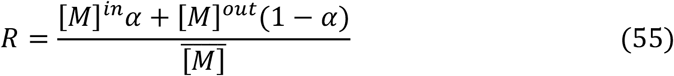

It’s easy to get:

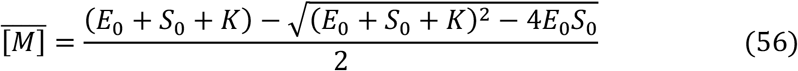

What’s more:

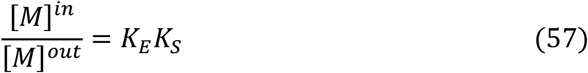

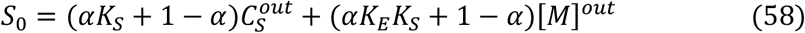

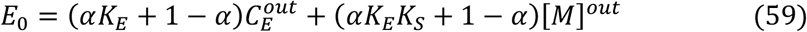

We then define:

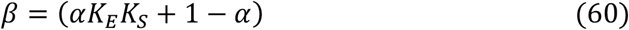

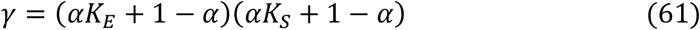

as a result:

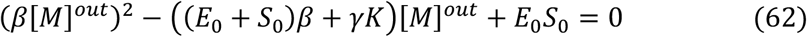

After solving the [*M*]^*out*^, we get:

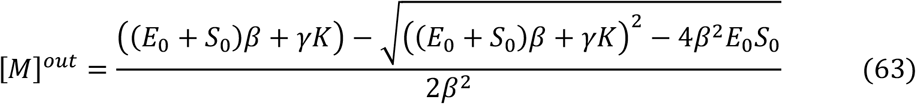

According to the equation (55),

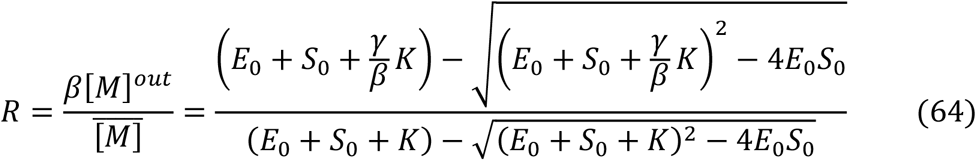

Where

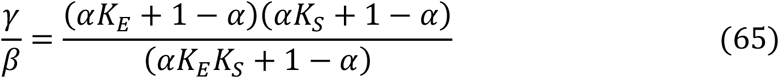

**Figure S1.**
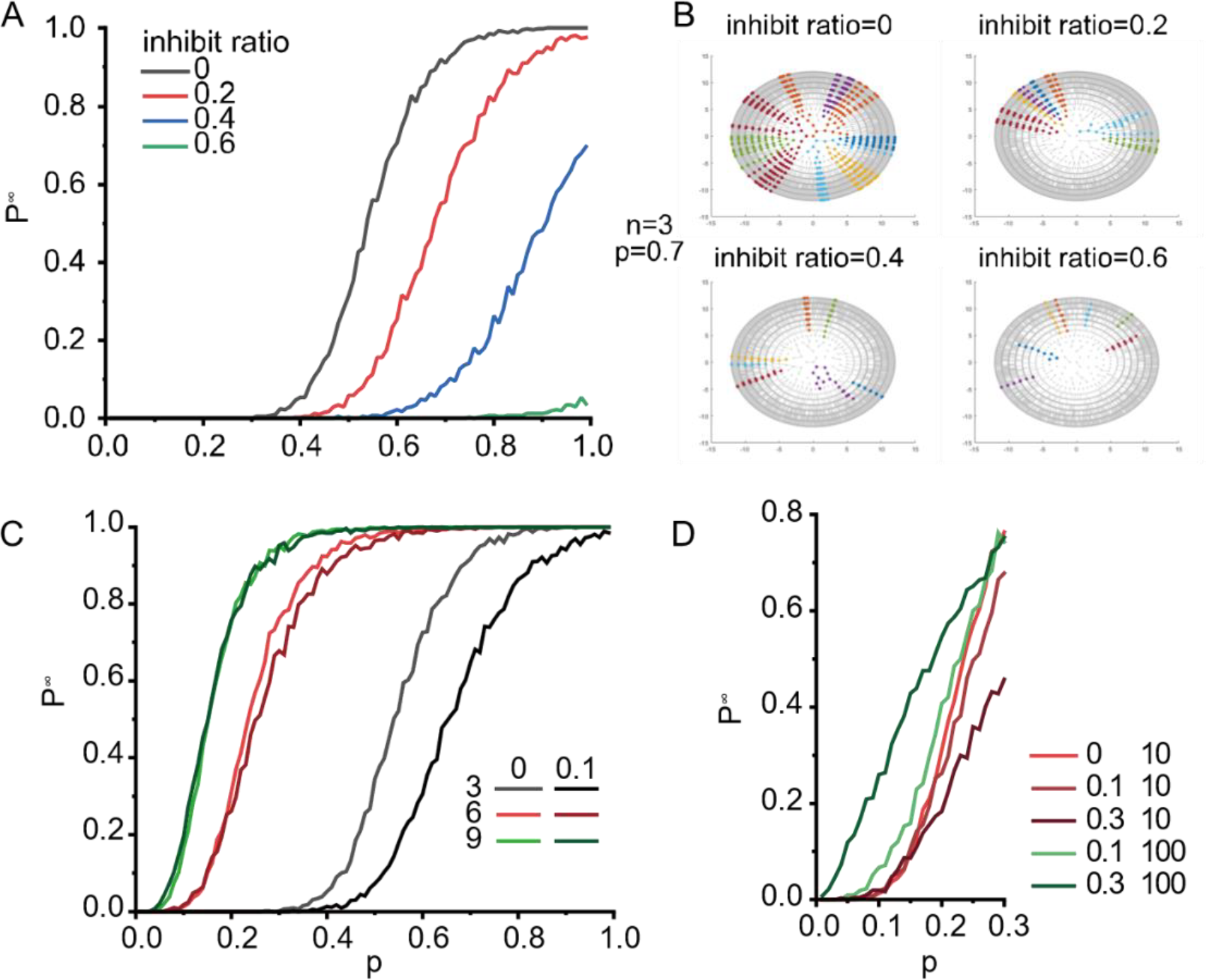
Simulations of the complex LLPS conditions. (A) The adding of inhibitors. (B) visualization of the results in A. (C) The synergy effect influences the percolation curves at different interaction valences. (D) The percolation curves of a hexavalent grid at synergy ratio of 0, 0.1 and 0.3 when the synergy gain was 10 or 100.

**Figure S2.**
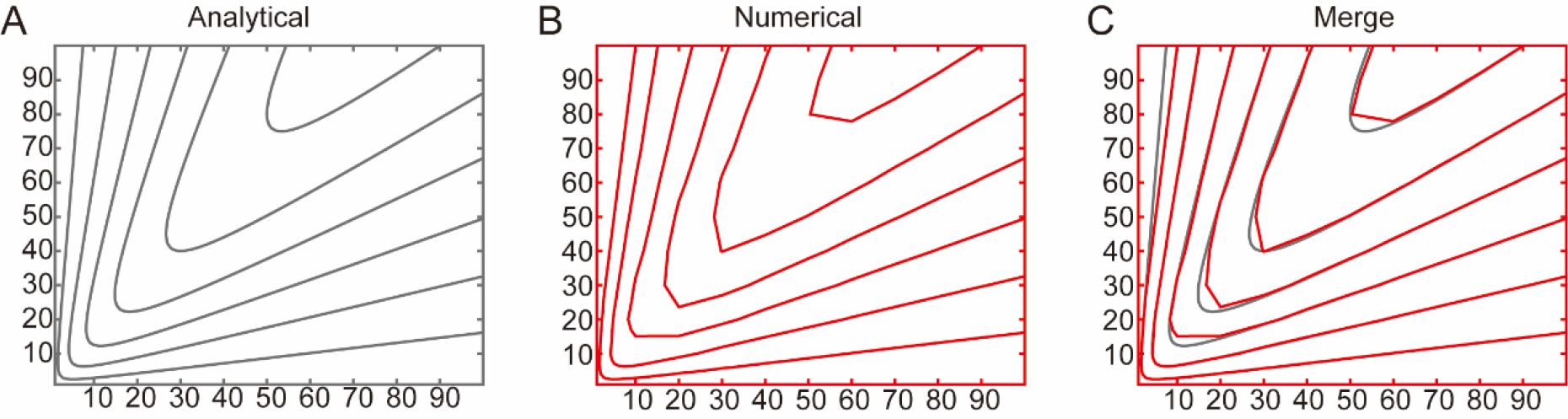
The overlap of the simulation and theoretical results of the “3+2” system.

**Figure S3.**
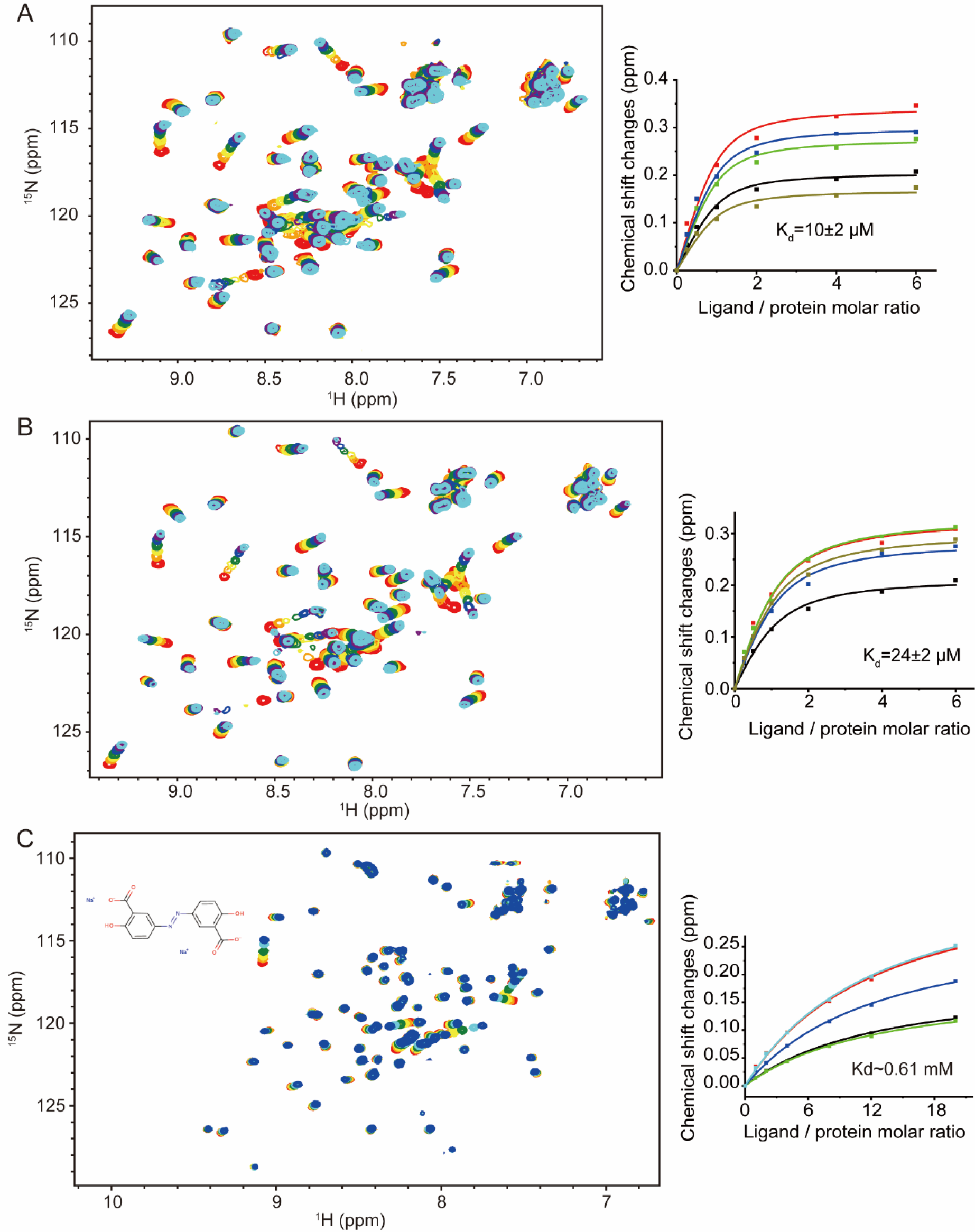
The results of NMR titrations. (A) the FCP1 peptide to T2FA, 150 mM NaCl. (B) the FCP1 peptide to T2FA, 225 mM NaCl. (A) the small molecule to T2FA, 150 mM NaCl.

**Figure S4.**
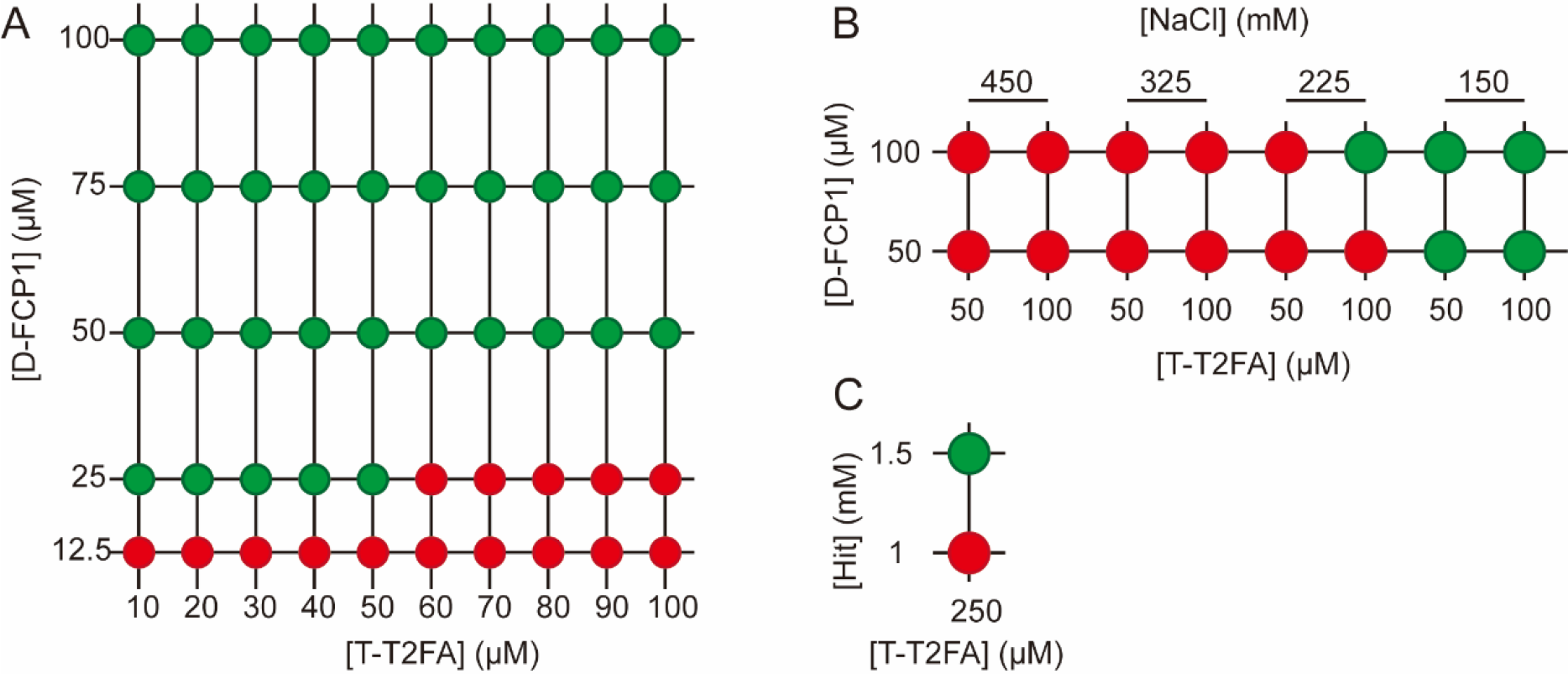
The LLPS results of the “3+2” system under different conditions. LLPS (green) or no LLPS (red) were showed by different colors.

**Figure S5.**
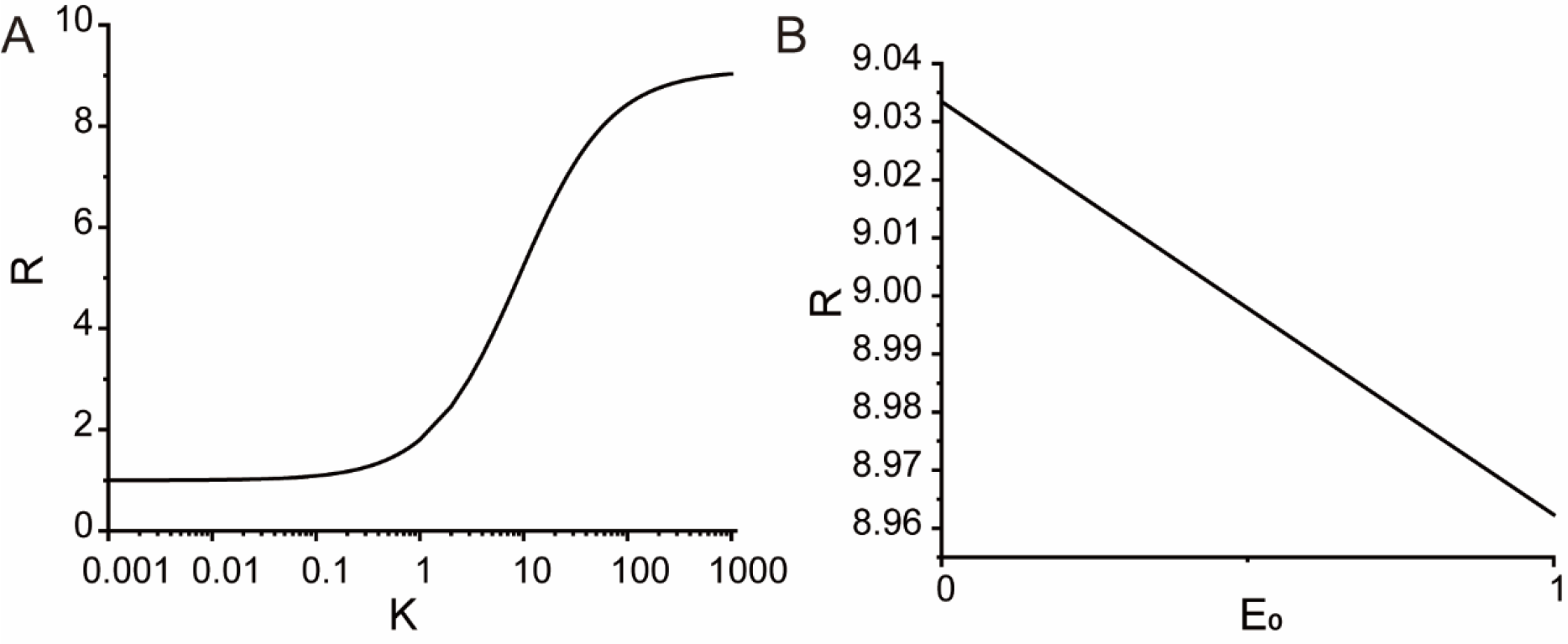
The reaction efficacy changed with the binding affinity and the concentration of the enzyme. (A) S_0_=1, E_0_=0.001, α= 0.001, K_E_=K_S_=100. (B) S_0_=1, K=1000, α= 0.001, K_E_=K_S_=100.

